# Examining the molecular clock hypothesis for the contemporary evolution of the rabies virus

**DOI:** 10.1101/2023.09.04.556169

**Authors:** Rowan Durrant, Christina A. Cobbold, Kirstyn Brunker, Kathryn Campbell, Jonathan Dushoff, Elaine A. Ferguson, Gurdeep Jaswant, Ahmed Lugelo, Kennedy Lushasi, Lwitiko Sikana, Katie Hampson

## Abstract

The molecular clock hypothesis assumes that mutations accumulate on an organism’s genome at a constant rate over time, but this assumption does not always hold true. While modelling approaches exist to accommodate deviations from a strict molecular clock, assumptions about rate variation may not fully represent the underlying evolutionary processes. There is considerable variability in rabies virus (RABV) incubation periods, ranging from days to over a year, during which viral replication may be reduced. This prompts the question of whether modelling RABV on a per infection generation basis might be more appropriate. We investigate how variable incubation periods affect root-to-tip divergence under per-unit time and per-generation models of mutation. Additionally, we assess how well these models represent root-to-tip divergence in time-stamped RABV sequences. We find that at low substitution rates (<1 substitution per genome per generation) divergence patterns between these models are difficult to distinguish, while above this threshold differences become apparent across a range of sampling rates. Using a Tanzanian RABV dataset, we calculate the mean substitution rate to be 0.17 substitutions per genome per generation. At RABV’s substitution rate, the per-generation substitution model is unlikely to represent rabies evolution substantially differently than the molecular clock model when examining contemporary outbreaks; over enough generations for any divergence to accumulate, extreme incubation periods average out. However, measuring substitution rates per-generation holds potential in applications such as inferring transmission trees and predicting lineage emergence.

**Author Summary:** Rabies is a neglected disease that kills around 60,000 people each year. After entering the body, the incubation period of the virus is usually less than one month, but can sometimes span months to years. While we normally assume a virus accumulates mutations at a constant rate, it is possible that rabies’ occasional long incubation periods mean that mutations accumulate at varying rates if the virus replicates (and thus mutates) more slowly during the incubation period. We compared how the rabies virus evolves over time using two simulation models where mutations either occur per unit time or per infection generation. We also calculated the mean substitution rate per infection generation, which can be useful for inferring linkage between related rabies cases. We found that at realistic substitution rates for the rabies virus, we could not distinguish between the two models. Our calculations show that in most generations no mutations are expected to occur. Thus, over a time period long enough to observe genetic divergence, occasional long incubation periods would be “cancelled out” by shorter than average incubation periods, meaning that the two models are almost equivalent. However our work suggests that modelling substitution rates per generation may be useful for epidemiological inference.

## Introduction

The molecular clock hypothesis assumes that the genomes of organisms accumulate neutral mutations at a constant rate over time, either across all lineages (the “strict molecular clock”) or within each individual lineage but with some degree of variation between them (clock models with this assumption include the relaxed and multirate clock models) (1–3). The ability to sample viral sequences through time, and the application of the molecular clock hypothesis to these sequences, has led to massive advances in using viral genetic data to investigate disease outbreaks (4). The clock rate, measured in substitutions per site per unit time, can be used to estimate how long ago pathogens diverged (5), and the date of infection of individual infected hosts (6). Combining the analysis of epidemiological and genetic data has allowed further insights into the history of outbreaks (7), and the introduction of geographic data provides estimates as to rates of spread and the frequency and source of introductions (8,9). However, in order to conduct these phylogenetic analyses, genetic divergence must increase appreciably over time in the dataset under investigation (10). Whether or not the viral population is measurably evolving, and thus whether it contains sufficient temporal signal for phylogenetic analysis, depends mainly on the evolutionary rate, the sequence length and the length of time sequences are sampled over being sufficiently high. Various methods exist to assess temporal signal, the most commonly used being root-to-tip divergence plots (11,12) implemented in tools such as TempEst (13), but these also include Bayesian evaluation of temporal signal (BETS) (14) and the date-randomisation test (15).

The rabies virus (RABV) is a negative-strand RNA virus, with a genome size of approximately 12 kilobases. While RNA viruses generally have high mutation rates due to a lack of proofreading by RNA polymerases, RABV has a substitution rate at the lower end of normal for single-stranded RNA viruses of between 1 x 10^−4^ and 5 x 10^−4^ substitutions per site per year (16–18). This may be due to strong purifying selection (16), or due to peculiarities of RABV. For example, the RABV genome is longer than average for RNA viruses, and genome length and evolutionary rate are negatively correlated (19), although this relationship appears to be weaker in single-stranded RNA viruses (20). A more unusual feature of RABV is that infections can exhibit extended incubation periods within the host. The median generation interval (the time between one individual becoming infected and then infecting another) is estimated to be 17.3 days in domestic dogs (21), with other studies estimating mean serial intervals of 26.3 days (22) and 45.0 days (23). Symptoms, infectivity, and death from rabies, however, can occasionally occur years after the initial infection event (24). The length of the incubation period is influenced by the route of exposure, with bites to the head and neck leading to more rapid disease progression than bites to lower extremities (25). RABV can remain in the muscle at the bite site for prolonged lengths of time before invading the host’s motor neurons and progressing through the nervous system, with limited, if any, infection of other muscle fibres (26). While some replication in the muscle cells has been observed (27), RABV replication at the inoculation site is not necessary for neural invasion (28). It is currently unknown precisely how the RABV replication rate in the host muscle cells and peripheral nervous system compares to the massive replication rate within the cells of the central nervous system and brain. However, work suggests that RABV replication in muscle cells may be reduced (29), and RABV replication in cultured rat sensory neurons may be 10- to 100-fold lower than replication rates in rat and mouse CNS neurons (30). Rabies infections that involve long incubation periods may, therefore, not lead to more accumulated mutations than those with shorter incubation periods, as viral mutation is strongly influenced by the replication process (31).

Changes in mutation rates through time due to long incubation periods may affect how we analyse RABV sequence data and interpret these analyses. A relaxed molecular clock is usually required to carry out phylogenetic analyses on rabies datasets, and it is not uncommon for there to be difficulties in applying these analyses due to “insufficient temporal signal”; usually referring to either no or a negative relationship between genetic divergence and time, or this relationship having a lot of noise and a very low R^2^ (32–36). RABV shows variation in substitution rate between lineages (18,37,38) which may be driven in part by differences in incubation periods. If the variable incubation period of rabies infections does cause deviation from the molecular clock model (exceeding the variation captured by relaxed or multirate clock models), this may negatively affect the accuracy of time-scaled phylogenetic trees and emergence date predictions. Conversely, if mutation does continue at a consistent rate during the incubation period, attention should be paid to extremely long incubators which could drive the emergence of new variants, as seen recently in chronic SARS-CoV-2 infections (39,40).

We hypothesised that reduced replication (and thus mutation) during the incubation period could cause rabies evolution to be better represented by a per-generation model of mutation than by the molecular clock model. We aim to clarify the nature of contemporary RABV evolution using in silico methods, comparing the root-to-tip divergence of sequences generated from synthetic outbreaks under per-unit time or per-generation mutation models, and comparing these to RABV genomic data from Tanzania. We also aim to calculate a per-generation substitution rate for RABV for future use as a parameter in transmission tree reconstruction algorithms.

## Methods

We investigate two contrasting mutational models for RABV – i.e., substitutions occurring on a per-generation vs. per-unit-time basis – using a simulation approach. We first generated synthetic RABV outbreaks using a branching process model (21) and then simulated these two mutation processes over the resulting transmission trees. From the synthetic sequences generated, we examined root-to-tip divergence and calculated variance explained (R^2^) from linear regressions, and compared these to the root-to-tip divergence of a set of RABV whole genome sequences from Tanzania. Finally, we developed a method to estimate the per-generation substitution rate for RABV and tested this on synthetic data before applying it to the Tanzanian RABV dataset.

### Rabies outbreak simulation

We simulated RABV mutation on branching-process simulations of rabies outbreaks. Outbreaks were simulated 100 times over a spatially explicit representation of Mara Region in northern Tanzania. In Serengeti District, where contact tracing data were available, the model was initialised with the three cases that occurred in the mean generation interval (*g*=27 days, based on contact tracing data) prior to 2017 (simulations were run over a dog population representing that in Mara region between 2017 and 2024). In the rest of Mara region, where there were no data to guide initialisation, we seeded with (0.01*Dg*)/365 cases, where *D* is the initial dog population in that area. If R_e_=1 (endemic transmission), this results in roughly 1% of the population becoming rabid over a year; contact tracing data suggest that incidence typically does not exceed that level (41). This led to a total of 273 initial cases in the region. Each case was assigned a number of offspring cases drawn from a negative binomial distribution (41) with mean (R_0_)=1.05 and dispersion parameter=1.33. The R_0_ value was chosen to result in a median number of cases each month that was roughly constant over time (over the 100 simulations), mimicking endemic disease. Movement of rabid dogs from their home locations to and between transmission locations followed a random walk with step lengths drawn from a Weibull distribution (shape=0.41, scale=0.13). We simulated occasional long-distance transport of dogs to a random location prior to their first transmission in 2% of cases (21). At each of a rabid dog’s transmission locations, another dog was randomly selected within the local 1km2 grid cell. If this dog was susceptible (i.e., not vaccinated or already incubating infection from a prior transmission event), rabies was transmitted. A generation interval was drawn for each new infection from a lognormal distribution (meanlog=2.96, sdlog=0.82), describing the time delay before it also became rabid and made its assigned transmissions. The step-length and generation-interval distributions were fitted using contact tracing data from Serengeti District, Tanzania (21). Branching process simulations were continued until 7 years had passed or rabies went extinct. Each synthetic case was assigned an individual ID, and for every case (except initial seed cases) we recorded the ID of the associated progenitor case. Dates of infection and transmission were recorded for each case.

We isolated complete transmission trees descending from each of the 273 initial cases from within one randomly selected synthetic outbreak. Transmission trees that contained over 100 cases (9 out of 273 trees in total, that ranged in size from 533 - 19,382 cases) were then used to generate synthetic sequence data. Across these trees, we see a mean generation interval of 26.6 days, and 2.5 and 97.5 percentiles of 3.90 and 94.11 days (Supplementary Figure S1). For each of the 9 trees the index case was assigned an initial 12kb genome sequence. Under the per-unit time mutation model, we determined the expected number of mutations by multiplying the substitution rate, the genome length and the length of the generation interval, for each case along the resulting transmission tree (because we assume mutations are neutral, the individual-level mutation rate is the same as the population-level substitution rate). The realised number of mutations was then drawn from a Poisson distribution, with this mean. We then randomly chose positions to change and new nucleotides to change them to. The resulting synthetic sequence data is referred to as the “time-based sequence data”. The generation-based model of mutation works as above, with the exception that the expected number of substitutions in a generation is constant and produces the synthetic “generation-based sequence data”.

### Divergence rate analysis

To investigate patterns of temporal divergence under the mutation models described above, we generated synthetic data with values of substitution rates ranging from 0.05 to 3 substitutions per genome per generation (or the per unit time substitution rate equivalent) and 4 population sampling regimes (from 1% of cases to 20%, informed by a previous study that estimated that routine surveillance for rabies rarely confirms more than 10% of circulating cases (42)). We calculated the genetic divergence as the number of nucleotide differences from the index case to each sampled case. For each of the nine transmission trees, we then compared genetic divergence with time under each scenario (substitution rate and sampling regime combination), using linear regression through the origin.

In order to compare our synthetic patterns of divergence over time to real rabies data, a root-to-tip divergence plot was also generated for a dataset of real RABV sequences (data from (43); Figure 1A) using TempEst (v1.5.3 (13)), with the best-fit root located (Figure 1B). These rabies cases occurred between 2001 and 2017 and were primarily from the Serengeti district and Pemba Island, with the remaining sequences from elsewhere in Tanzania (Figure 1A inset). Sequence acquisition and tree building methods are detailed in (43).

**Fig 1.**
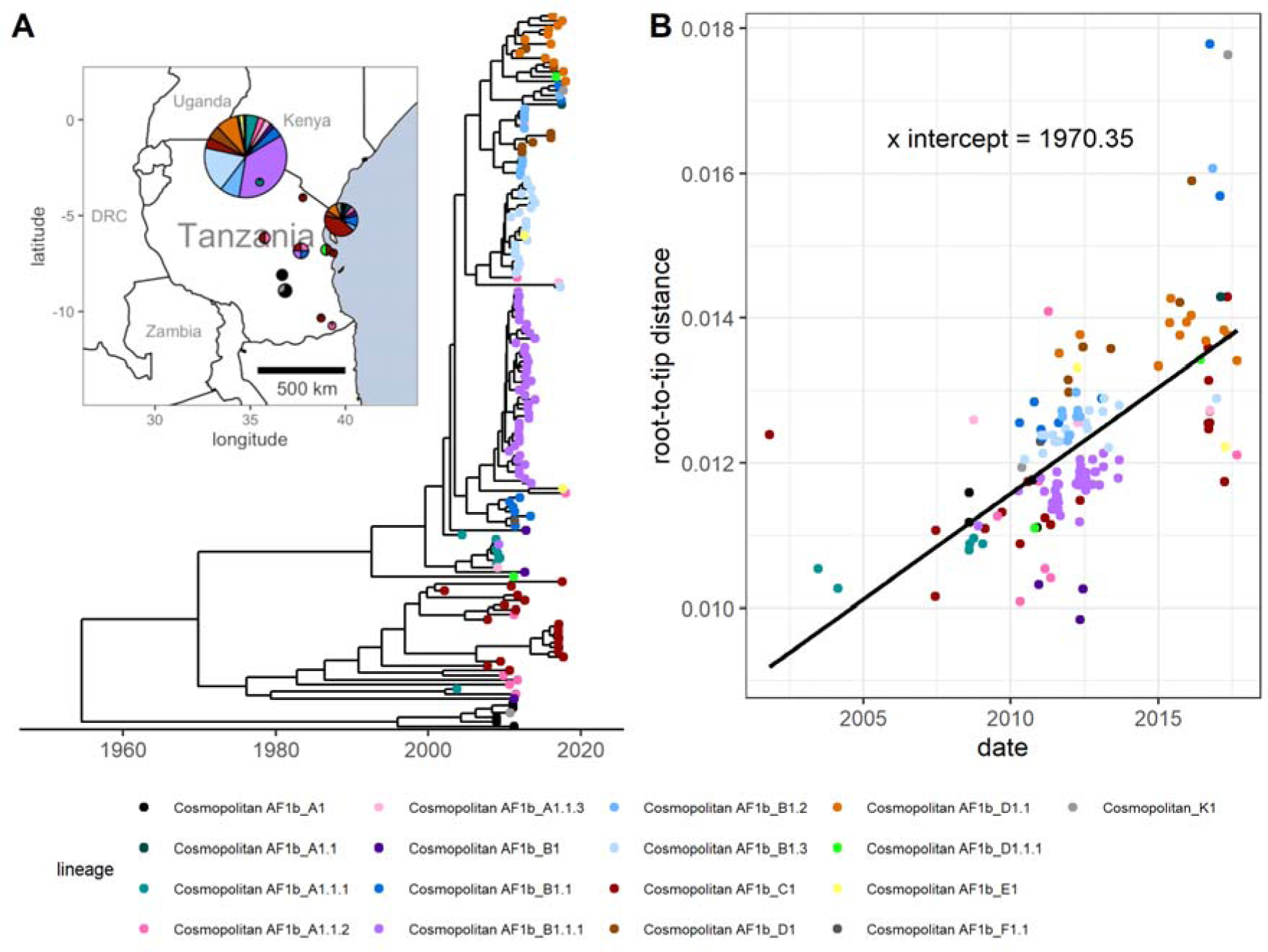
Phylogeny of real rabies virus whole genome sequences from Tanzania and root-to-tip divergence. **(A)** The time-scaled tree (43) used to generate the root-to-tip divergence plot and to calculate the per-generation substitution rate. The inset map shows the approximate locations that the samples were collected from, and the lineages present in each location. Map point size represents the number of sequences in this dataset from district centroid locations. Base map data is from Natural Earth (naturalearthdata.com), via the *maps* R package. **(B)** The corresponding root-to-tip divergence plot. Point colours represent RABV lineage.

### Calculating the per-generation substitution rate

We updated a method of calculating the per-generation substitution rate previously used in eukaryotes (44) by using Bayesian posterior estimates of the clock rate and the generation interval. We assessed this method’s accuracy using the synthetic outbreak sequence data, before applying it to the aforementioned set of RABV whole genome sequences.

To estimate the mean per-generation substitution rate, we analysed sequence data with BEAST, and multiplied the posterior rate estimate for each MCMC sample (excluding the burn-in period) by the generation-interval lengths sampled from the posterior of a simple Bayesian analysis and then multiplied again by the genome length. The mean and 95% credible interval of the estimate of the per-generation substitution rate for the RABV dataset was calculated by taking the mean and the 2.5% and 97.5% percentiles of the resulting multiplied posteriors.

To evaluate the accuracy of this method in estimating the mean per-generation substitution rate, we also applied it to synthetic sequence data generated from outbreaks using the per-generation mutation model as described above, under different substitution rates (11 values ranging from 0.05 substitutions per generation to 3 substitutions per generation) and case sampling rates (1%, 5%, 10% or 20% of cases sampled) across the 9 transmission trees that contained at least 100 cases. Subsampled synthetic datasets containing more than 2000 sequences were not analysed as this number exceeds the total whole-genome RABV sequences currently available on the RABV-GLUE database (45), and is unrealistic in the context of examining individual rabies outbreaks. BEAST log files were generated from these sequences using BEASTGen version 1.0.2 and BEAST version 1.10.4 (46). We chose to use a JC substitution model with a strict clock, no site heterogeneity due to our per-generation mutation model used in the simulations having equal probability of any site or base being chosen and assumed constant population size. We used a tracelog frequency of 1000 and a sufficiently long chain length for the effective sample size (ESS) of each parameter to exceed 200 when analysed using Tracer (47), and a 10% burn-in period. We applied the substitution rate calculation method to these phylogenetic trees, and assessed the accuracy of the resulting mean per-generation substitution rates by comparing them to the parameter values used to generate the synthetic sequences, using the natural log of the ratio:

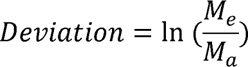

where M_e_ is the mean estimated per-generation substitution rate and M [is the actual substitution rate, where a deviation of zero means perfect accuracy.

The same method was applied to the dataset of 153 RABV sequences sampled from across Tanzania (data from (43); Figure 1A). The mean per-generation substitution rate was calculated, and distributions were fitted from the multiplied generation interval and clock rate posteriors (generation interval posteriors based on values from (21) for the Tanzanian dataset, extracted directly from the lognormal distribution used in simulations, and clock rate posteriors taken from the BEAST log file of the time-scaled tree from Lushasi et al. (43)) and genome length as described above. We compared different distributions (Gamma and Lognormal) for estimating substitution rates and selected the best fitting distribution by AIC. We also calculated the probabilities of between 0 and 10 SNP differences occurring across 1, 5 or 10 infection generations. For this calculation we simulated mutations arising at a Poisson rate with lambda drawn from the fitted substitution rate distribution. The means and 95% confidence intervals were calculated from the 10,000 simulations.

### Data and code availability

All code is available at https://github.com/RowanDurrant/Rabies-Mutation. Analyses were conducted using the R programming language (48). The beta regression curve and prediction interval in Figure 2C was generated using the ‘betareg’ R package (49). RABV lineages were assigned using MADDOG (45).

**Fig. 2:**
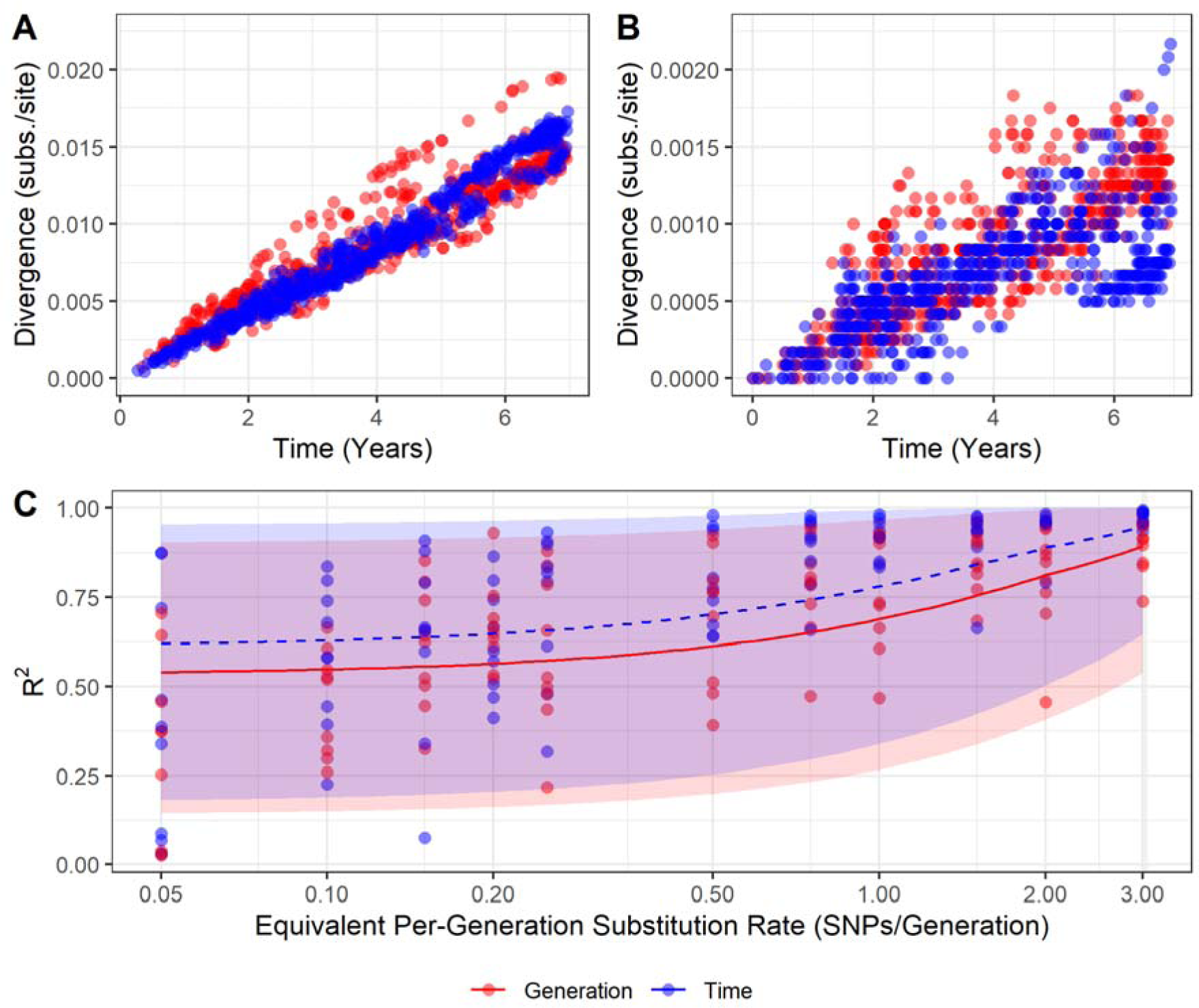
Temporal genetic divergence varies under two models of mutation. **(A)** Root-to-tip divergence plots for synthetic sequences produced using time-based and generation-based mutation models, equivalent to 2 substitutions per genome per generation and **(B)** equivalent to 0.2 substitutions per genome per generation. Note that the y-axis scales differ by an order of magnitude between A and B. These data are from running mutation models over the same single transmission tree and have a case sampling rate of 5% (i.e., 621 cases sampled of 12,434 total). **(C)** The R^2^ values obtained from regression through the origin of root-to-tip divergence of synthetic data from the time-based and generation-based models. Point colour indicates the mutation model used to generate the data. Lines represent beta regressions with logit links fit to data points, and shading represents the 95% prediction interval. The X axis is log scaled. 5% of cases were sampled here; sampling rate had little effect on R^2^ (Supplementary Figure S2).

## Results

### Root-to-tip divergence analysis

At higher per-generation substitution rates (1 substitution per genome per generation and above), distinct differences can be seen between root-to-tip divergence plots from the two models of mutation (Figure 2A). The synthetic data generated from the per-generation mutation model shows “stray” clusters or ridges of points both above and below the main funnel of points, illustrated in the example in Figure 2A. Divergence plots from synthetic data generated from the time-based model of mutation have less variance and do not exhibit this pattern. At lower substitution rates (below 1 substitution per generation), no such pattern is clearly distinguishable (Figure 2B). When the cases represented by the high-divergence points from the per-generation model in Fig. 2A are visualised in a transmission tree, they are mainly confined to a single chain (Supplementary Figure S1).

Root-to-tip divergence plots derived from synthetic transmission trees using the time-based mutation model had, on average, higher R^2^ values than those from synthetic transmission trees using the per-generation mutation model, although this is more difficult to distinguish below a substitution rate of 0.5 substitutions per genome per generation (Figure 2C). As the substitution rate increases, the R^2^ values across both mutation models increase. The case sampling rate appears to have little effect on R^2^ (Supplementary Figure S2).

The root-to-tip divergence plot of the Tanzanian RABV dataset more closely resembles those of lower substitution rate simulations, where it is difficult to determine any difference between the models of mutation (Figure 1). While most lineages surround the regression line, some (for example, Cosmopolitan AF1b_B1) group below the line, but without forming a distinguishable “ridge”.

### Substitution rate calculation

The accuracy of our method used to calculate per-generation substitution rate remains similar at all but the lowest values of substitution rate (Figure 3), with a tendency to underestimate the substitution rate (meaning that the estimated substitution rate is below the substitution rate parameter used to generate the synthetic data; mean natural log of the ratio of −0.18 and root-mean-square of 0.54, where values of 0 indicate perfect estimates). Accuracy appears to be more influenced by the number of sequences used in the BEAST analysis than by the case sampling rate itself; the mean natural log of the ratio falls to −0.36 when fewer than 50 sequences are used (root-square-mean of 0.95).

**Fig. 3:**
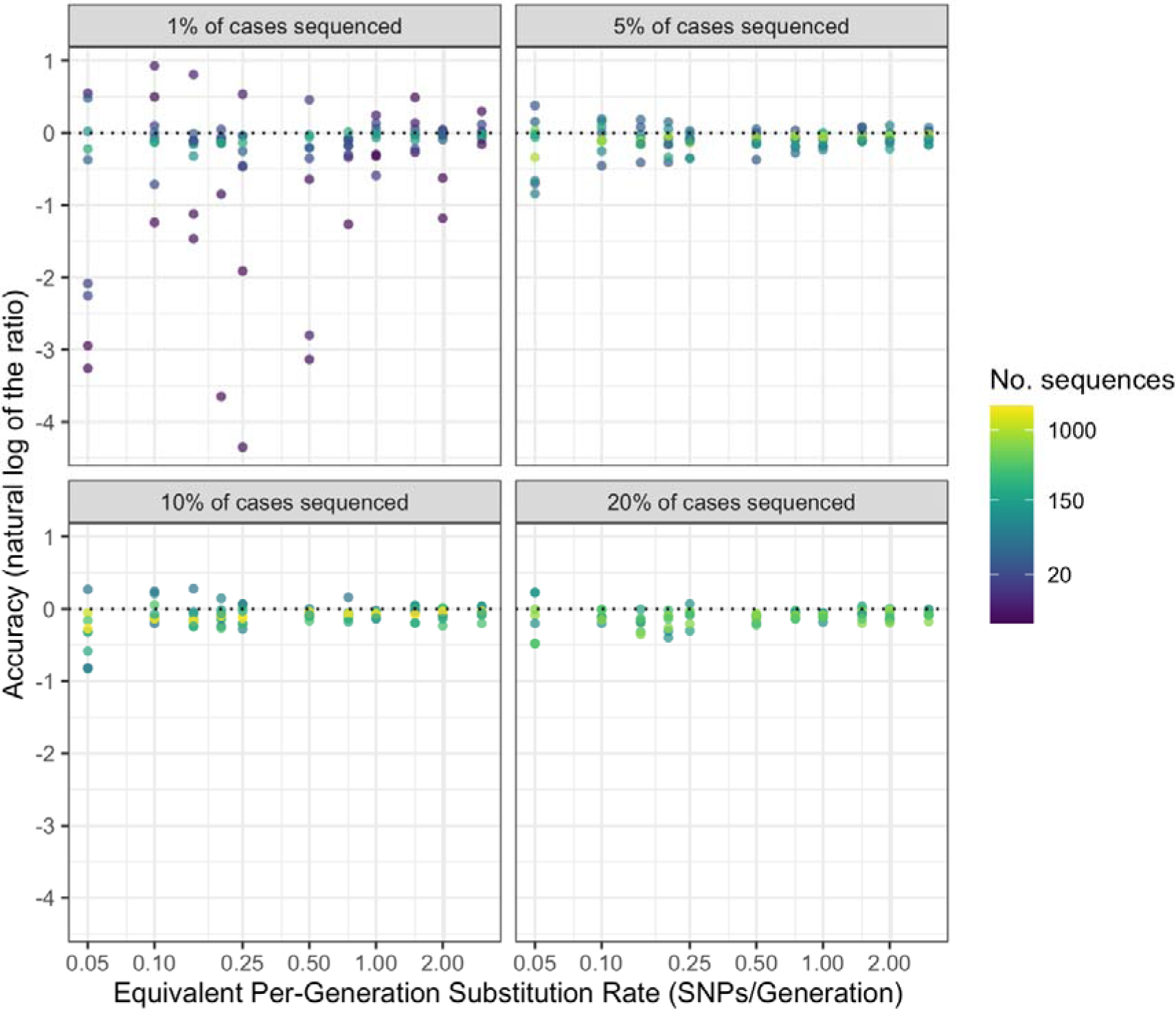
Accuracy of per-generation substitution rate predictions for different numbers of sequences, substitution rates and sampling rates. Facets indicate case sampling rate. The dotted line represents perfect accuracy. X axis and colour scale are log transformed.

The Tanzanian RABV dataset from which we estimated the per-generation substitution rate contains 153 sequences in total, and the accompanying time-scaled phylogenetic tree has a root-to-tip height of approximately 65 years, although the sequences spanned just 16 years as they were sampled from 2001 to 2017 (with 46.7% from years 2011-2012). These sequences were largely complete; 98% of sequences were >95% complete (>11,327 kb in length). The mean per generation substitution rate of RABV in this dataset was estimated to be 0.171 (95% credible interval: 0.127 - 0.219). The best fitting distribution by AIC to the output of the multiplied Bayesian posteriors was a Gamma distribution with the parameters shape = 51.69 and rate = 301.8.

Using the calculated per generation per genome substitution rates, we calculated the probability of different numbers of substitutions occurring over 1, 5 and 10 generations, drawing the per-generation substitution rate (λ) from the above distribution (Figure 4). Over many generations it is still quite likely for zero substitutions to occur; after 10 generations, the probability of zero substitutions having occurred is 0.19.

**Fig. 4:**
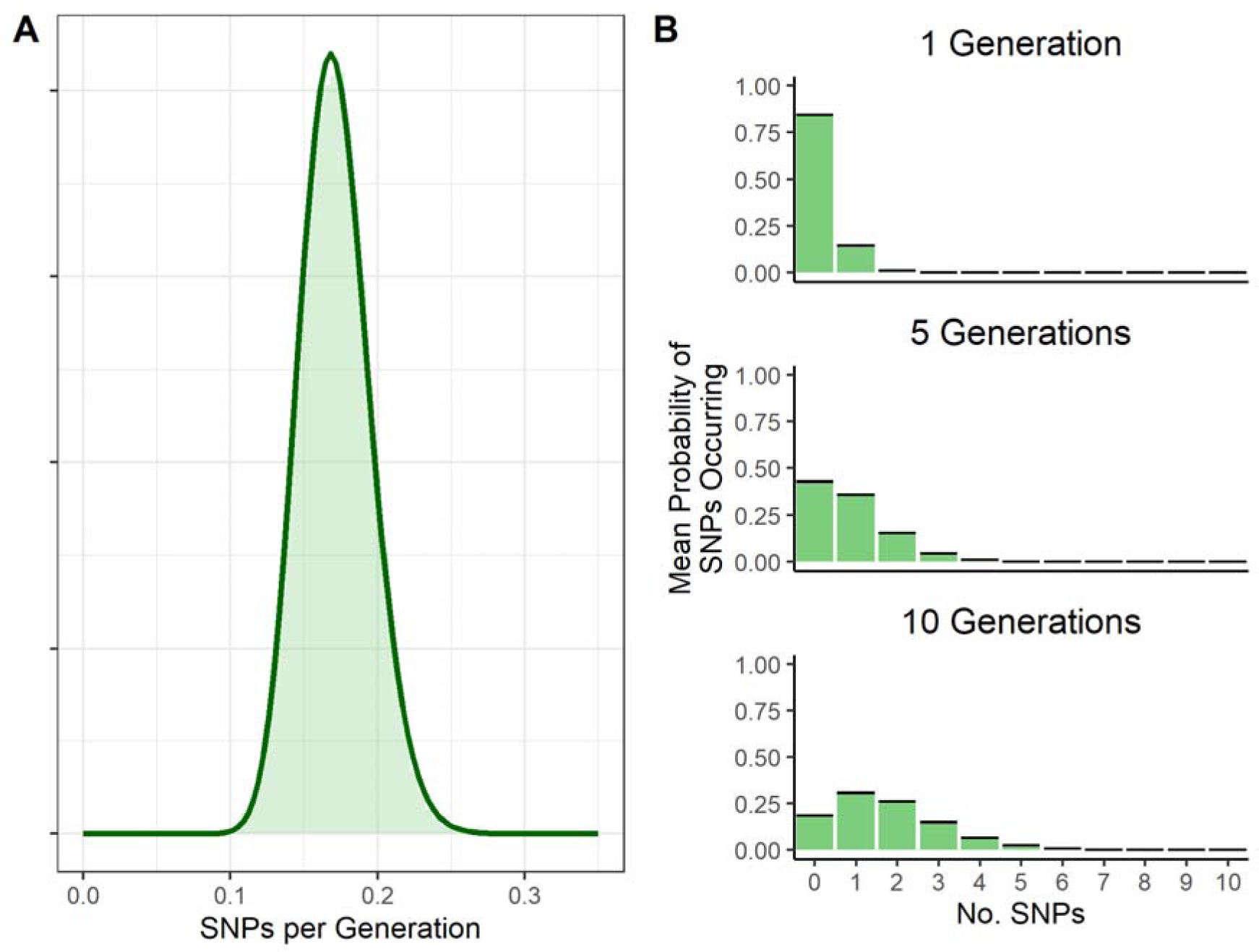
Probability distributions of the mean per-generation substitution rate and substitutions occurring over generations. **(A)** estimated probability distribution of the per genome per generation substitution rate from Tanzanian RABV sequences, with underlying histogram of multiplied Bayesian posteriors of clock rate and generation interval. **(B)** probability distribution of SNPs occurring over 1, 5 and 10 generations. The λ value for a Poisson rate of SNP occurrence is drawn from the SNPs per generation distribution fitted in Figure 4A. Black bars represent the 95% confidence intervals (which are very tight).

## Discussion

It can be difficult to get sufficient temporal signal for RABV sequence datasets, which we hypothesised could be due in part to its variable incubation periods. We hypothesised that a per-generation model of mutation may be more representative of RABV evolution than a purely time-based model. We found that substantial differences in root-to-tip divergence patterns between synthetic outbreaks using generation-based and time-based models of mutation could be observed only at high underlying substitution rates. The substitution rate for the Tanzanian RABV sequences examined (∼0.17 substitutions per genome per generation) was in the range where divergence patterns in the two models were extremely similar. We can thus assume that the two models will give extremely similar results on the relevant time scale. As we observed increasing divergence over time with reasonable R^2^ values within this substitution rate range, it implies that variable incubation periods alone do not fully account for the challenge in obtaining temporal signal. Therefore, other factors such as insufficiently long sampling windows for the substitution rate are likely to be responsible (50). This is an important consideration for analysing RABV sequences from new outbreaks, or from endemic areas where sampling is opportunistic. As RABV has a substitution rate lower than many other RNA viruses, longer sampling windows are required to achieve a sufficient temporal signal.

The observation of little difference between root-to-tip divergence plots derived from the two mutation models at substitution rates below 1 substitution per genome per generation is likely because of averaging; multiple generations of infection are expected to have occurred per substitution that arises on the viral genome. Over the many generations needed before significant levels of viral genetic diversity are reached, the influence of any unusually long incubation periods will be damped by the opposite influence of unusually short incubation periods, eventually becoming indistinguishable from clock-like behaviour. On the other hand, at higher substitution rates ridges form on the root-to-tip divergence plots under the per-generation model of mutation but not under the per unit time model. While not affecting the overall clock rate, these ridges reduce the overall R^2^, and may be better analysed using a separate local clock (51). The cases in these ridges almost all descend from a common ancestor (Supplementary Figure S1), suggesting that a single unusually long or short incubation period can affect which phylogenetic analyses we perform. Ridges caused by these incubation periods can be distinguished from ridges caused by rate variation between lineages as they will be parallel to the main cluster of points in the plot, whereas points belonging to lineages with a different substitution rate will have a different slope. Studies examining the number of substitutions occurring between successive sequenced cases, and whether this increases when the secondary case’s incubation period is unusually long, could clarify the exact relationship between substitutions, generations, and time. More detailed data will be required to investigate this further.

We calculated RABV’s mean per-generation substitution rate to be approximately 0.17 substitutions per genome per transmission generation. This estimate is lower than those for other RNA viruses, such as SARS (2 substitutions per genome per human passage (52)), SARS-CoV-2 (0.52 substitutions per genome per 5.8-day generation interval (53)) and Ebola virus (0.875 substitutions per genome per 14-day generation interval (54)). RNA viruses that undergo periods of reduced replication or complete latency often show reduced substitution rates, with one extreme example being HTLV-1/2 (55,56). However, we would not expect this to affect the per-generation rate. The low per-generation substitution rate seen in rabies is therefore likely due to mutation being constrained by other factors, such as strong purifying selection (16), and likely contributes to the difficulties in obtaining sufficient temporal signal for phylogenetic analyses. Previous studies suggest that for viruses in this substitution rate range, sampling windows of up to 30 years may be required to overcome the phylodynamic threshold (15); for comparison, SARS-CoV-2 achieved sufficient temporal signal within two months of the start of the pandemic (50).

We can predict from the estimated per-generation substitution rate that identical sequences are likely to have less than 5 intermediate generations between them (probability of fewer than five generations occurring before a mutation occurs > 0.49 by repeated sampling of a Poisson distribution with a lambda of 0.17), but still have a non-negligible probability of being more distantly related. While the low substitution rate means that comparing the number of SNPs between sequences alone may not be an effective method of determining infector-infectee relationships, it could be used in conjunction with temporal and location data to make more accurate predictions of transmission events by ruling out relationships between more distantly related transmission chains co-circulating in the same area, as in (57). Notably, our Poisson distribution of the number of substitutions occurring in one generation is visually very similar to the genetic signature distribution reported in Cori *et al*. ((57), Fig S1), despite different methods and RABV datasets being used in their calculations. It is likely, however, that our estimate of the per-generation substitution rate is lower than the mean number of SNPs expected between sequences from a primary and secondary case, due to the time-based substitution rate being affected by purifying selection (58). Further analysis comparing the estimated per-generation substitution rate to realised SNP distances between primary-secondary case pairs could quantify this difference.

While the Jukes-Cantor model was the most appropriate to use on our synthetic data due to the simplicity of the mutation models, phylogenetic analyses on real RABV genomes usually use a more complex model, such as the GTR + G substitution model used to generate the Tanzanian tree shown in this study (43). This, along with the simplicity of our mutation model as well as sampling biases in the real dataset, may affect how comparable synthetic root-to-tip divergence plots are to the real data.

While the molecular clock has proven critical for gaining insights into the history and dynamics of disease outbreaks, the epidemiological characteristics of a virus should be considered when choosing how to measure viral evolution. In this study, we determine that the per-generation model is not likely to produce substantially different results from the molecular clock model when analysing contemporary RABV evolution. We also estimate the mean per-generation substitution rate of RABV for future use in transmission tree reconstruction and efforts to estimate outbreak sizes and lineage emergence rates. Given that many different lineages circulating simultaneously is seemingly a common occurrence in areas with endemic rabies, it is important to investigate whether these lineages vary in evolutionary rate and generation interval length, and ascertain the potential effects on phylogenetic analyses.

## Supporting information captions

**Supp. Fig. S1: histogram of generation intervals from the simulated outbreaks.** Vertical dashed lines represent the median (blue) and mean (red) generation interval.

**Supp. Fig. S2: points in the offshoot ridge predominantly occur in one transmission tree. (A)** root-to-tip divergence plot (2 SNPs/genome/generation, 5% of cases sequenced) with offshoot ridge points highlighted in red. Offshoot ridge points are defined in this plot as having a divergence rate above 8×10^−6^ substitutions/day and occurring after day 750. **(B)** transmission tree of the simulated outbreak with offshoot ridge cases highlighted in red. Graph edge length is not proportional to time or divergence.

**Supp. Fig. S3: Sampling rate does not impact the R^2^ of root-to-tip divergence plots from synthetic data.** Plot is faceted by the proportion of the total number of cases in the outbreak sequenced, point colour represents mutation model.

