## Supplementary information for "Examining the molecular clock hypothesis for the contemporary evolution of the rabies virus"

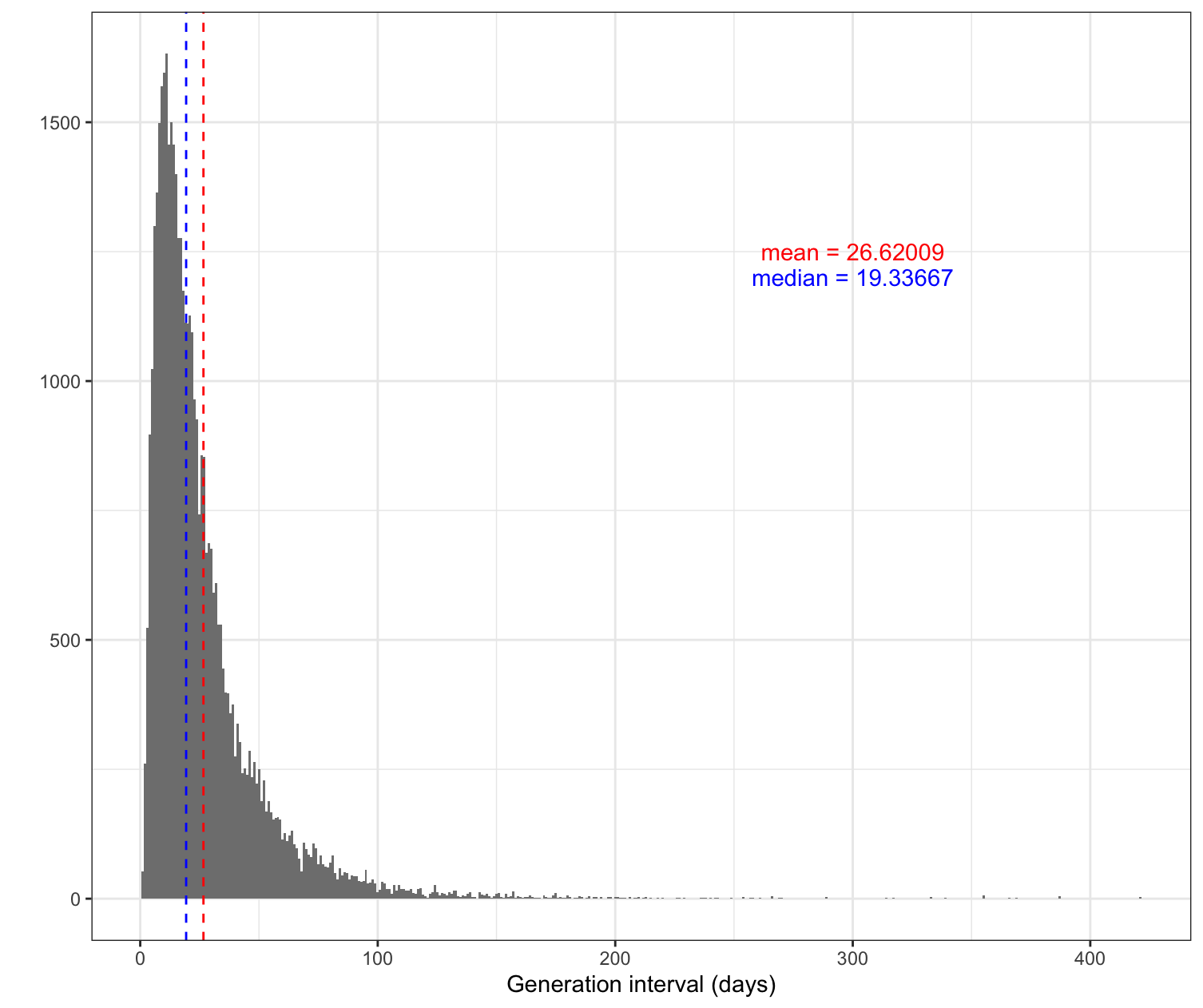

**Supp. Fig. S1: histogram of generation intervals from the simulated outbreaks.** Vertical dashed lines represent the median (blue) and mean (red) generation interval.

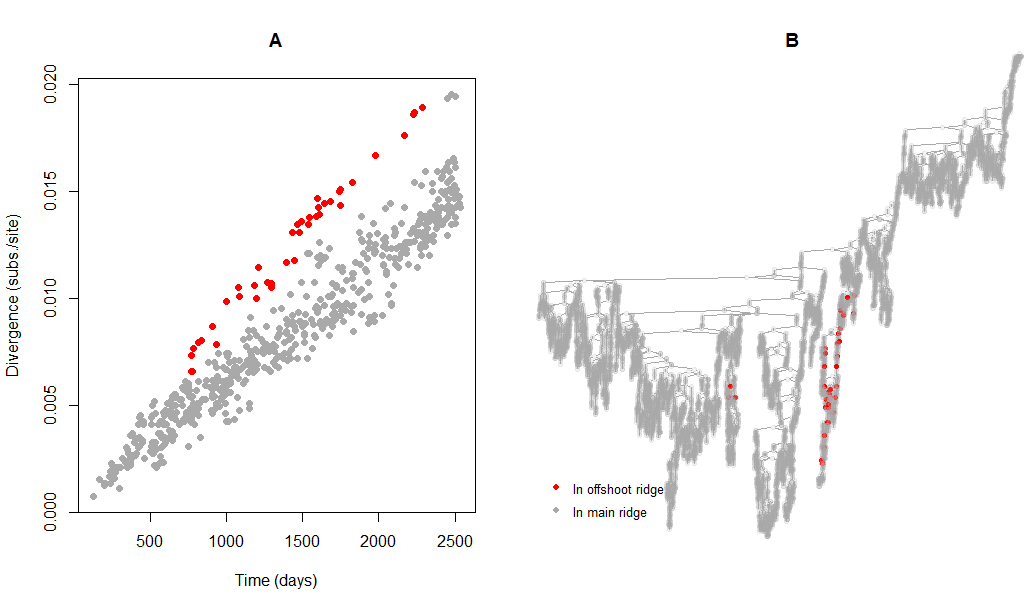

**Supp. Fig. S2: points in the offshoot ridge predominantly occur in one transmission tree. (A)** root-to-tip divergence plot (2 SNPs/genome/generation, 5% sampling rate) with offshoot ridge points highlighted in red. Offshoot ridge points are defined in this plot as having a divergence rate above 8x10^-6^ substitutions/day and occurring after day 750. **(B)** transmission tree of the simulated outbreak with offshoot ridge cases highlighted in red. Graph edge length is not proportional to time or divergence.

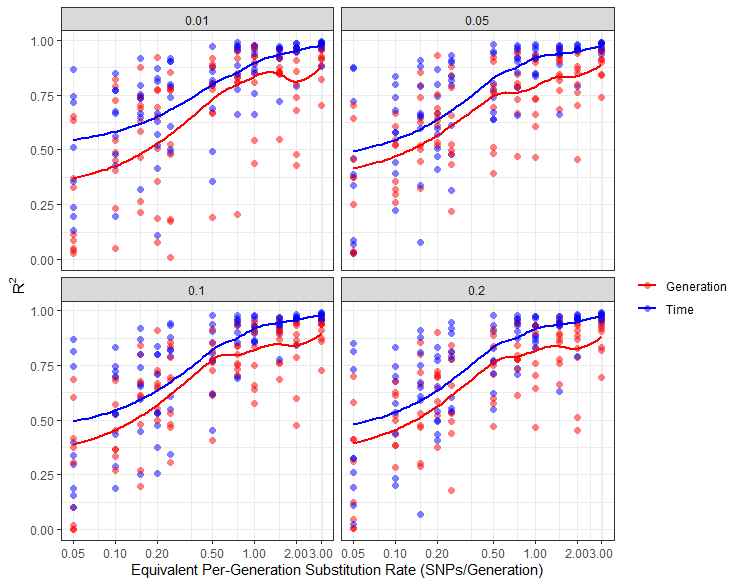

**Supp. Fig. S3: Sampling rate does not impact the R^2^ of root-to-tip divergence plots from synthetic data.** Plot is faceted by sampling rate as a proportion of the total number of cases in the outbreak, point colour represents mutation model.
